# Social relationships promote access to food and information in wild jackdaws

**DOI:** 10.1101/2025.10.25.684536

**Authors:** Luca G. Hahn, Guillam E. McIvor, Ines Fürtbauer, Andrew J. King, Alex Thornton

## Abstract

Many social animals form differentiated social relationships that can influence fitness, yet the mechanisms behind these benefits are often unclear. Does the presence of social partners during foraging enhance access to food or information? Might individuals also incur short-term costs, for example by foregoing feeding opportunities to remain with and support social partners? Here, we investigated how social relationships shape foraging decisions and outcomes in wild jackdaws *(Corvus monedula)* using automated *RFID* feeding stations where two birds could perch, but only one could feed with the other queueing behind (“dyadic events”). Bonded social partners (pair-bonded mates and direct kin) engaged in these dyadic events more frequently and coordinated their visits, as shown by shorter dyadic arrival latencies. Bonded partners also showed higher social tolerance, with feeding individuals benefitting from increased food intake when their partner was present, while queuing individuals accepted short-term opportunity costs to remain with them. For juveniles, queuing with parents promoted social learning about feeder use. These findings show that social relationships influence decision-making and foraging performance in the wild, revealing both benefits and costs of maintaining close bonds. Short-term effects on foraging success may therefore contribute to long-term fitness advantages of strong social relationships.

## Introduction

Many animals form and maintain differentiated social relationships among both kin and non-kin (1,2). Such social relationships, typically defined as repeated affiliative social interactions between specific individuals (1,3), result in structured social networks (4–6) and may provide important benefits for individuals. For instance, correlational studies on ungulates (7,8), primates (9,10), and cetaceans (11,12) indicate that individuals with stronger social relationships, or those who are more central in the wider social network, tend to be healthier, have higher reproductive success, and live longer (13–19). However, the mechanisms through which these benefits are obtained are generally unclear (20,21). To understand how these benefits (and the associated costs) arise, we need to investigate the immediate consequences of individuals’ decisions about whom to associate with and how to behave towards others. For instance, the short-term consequences of individuals’ decisions when foraging with others could provide a mechanism through which long-term benefits of social relationships emerge.

During foraging, animals may benefit from preferentially associating and coordinating activities with bonded partners, such as pair-bonded mates, kin, or “friends”. Individuals choose to forage alongside bonded partners in various species, including dwarf mongooses *(Helogale parvula)* (22), vampire bats *(Desmodus rotundus)* (23), Assamese macaques *(Macaca assamensis)* (24), and chacma baboons *(Papio ursinus)* (25). To achieve foraging benefits, bonded partners need to coordinate their foraging and show social tolerance (i.e. proximity without aggressive behaviour). This requires decision rules to discriminate among individuals: for example, domestic dogs *(Canis lupus familiaris)*, wolves *(Canis lupus)* (26), and vervet monkeys *(Chlorocebus pygerhythrus)* (27) show higher social tolerance towards bonded partners than other individuals. Social tolerance between bonded partners may also enable better access to contested resources and social information, potentially resulting in increased immediate food intake (28) and social learning (29–31) which could be particularly important for younger, naïve individuals. For instance, in common ravens *(Corvus corax)* (32), Barbary macaques *(Macaca sylvanus)* (33), and bearded capuchins *(Sapajus libidinosus)* (34), proximity to individuals solving experimental extractive foraging tasks, requiring social tolerance, predicted social learning by observers.

To understand the role of social relationships and social tolerance during foraging, we must consider not only the benefits, but also the costs. For example, individuals may trade longer-term benefits of maintaining group cohesion (e.g. reducing predation risk) with shorter-term costs (e.g. reduced foraging success). Foraging experiments with chacma baboons show this to be the case: lower-ranking baboons follow the movement decisions of higher-ranked baboons to food patches where they will receive less food but remain as a cohesive troop (35). In other cases, costs may be linked to specific dyadic social bonds. Great tits *(Parus major)*, for example, reduce their own foraging intake to remain near their foraging pair-bonded partner (36). This could reflect obligate bi-parental care, as individuals may be willing to incur short-term costs to ensure that their partner has sufficient resources to contribute to rearing offspring. This might be especially important for genetically monogamous species, where the fitness outcomes of both partners are highly interdependent, but this has yet to be examined.

An ideal system to investigate social decision-making during foraging is the jackdaw *(Corvus monedula)*, a highly social member of the Corvidae family. Jackdaw society is centred around long-term, genetically monogamous pair-bonds, which can last a lifetime (37). Pair members’ activities are coordinated throughout the year (38) and across different contexts, including nest building (39), incubation (40), flocking (41), and foraging (42). Another important social relationship in jackdaws is that between first-order kin, particularly between parents and their offspring. Juveniles remain dependent on, and retain close associations with, their parents as they transition to independence in the weeks and months after fledging (43). These strongly bonded partners are embedded in a dynamic social network with fluid membership (44), which means individuals regularly interact with others with whom they share varying levels of familiarity and relationship strength. Therefore, while individuals may form enduring social associations with familiar individuals that are not pair-bonded mates or kin (i.e., “friends” (2)), many interactions with less familiar individuals may be characterised by competition and limited social tolerance. Jackdaws are generalist foragers that regularly forage socially (43), at monopolisable food sources (43), facilitated by social information use (45,46). We therefore presented *RFID* feeding stations such that one jackdaw could feed at a time, while another could queue behind at a second perch in close proximity (hereafter referred to as “dyadic events”). This allowed us to record the identity, proximity, and feeding behaviour of individuals, providing detailed data on social interactions during foraging.

We tested the overarching hypothesis that jackdaw social relationships shape decisions and interactions at feeding stations, with consequences for the costs and benefits of accessing food and information. We compare strong relationships (bonded partners: pair-bonds and direct kin) to other, more transient relationships and test several linked predictions: *(1) Social bias*. Bonded partners should visit dyadic feeding stations together more frequently compared to other dyads. *(2) Coordination*. We predicted *(a)* bonded partners should arrive at the feeding stations closer in time than other dyads, and *(b)* the latency of arrival should be repeatable, reflecting coordinated foraging over time. *(3) Social tolerance*. We predicted *(a)* that bonded partners should queue for longer compared to other dyads, with queuing partners incurring a short-term opportunity cost, and *(b)* the bird at the feeder perch benefitting from a longer feeding duration when accompanied by a bonded partner. As above, we also expected that *(c)* the duration of dyadic events should be repeatable for dyads. *(4) Social learning*. We predicted *(a)* that juveniles that engaged in dyadic events with their parents should visit feeding stations earlier and more often than other juveniles that did not engage in dyadic events with their parents. *(b)* We predicted that naïve juveniles queueing behind juveniles that had previously followed their parents would visit feeders more often than those naïve juveniles who did not queue behind knowledgeable peers.

## Methods

### Study system

We studied jackdaws at two field sites in Cornwall, United Kingdom: site *Y*, located in a churchyard and surrounding fields in the village of Stithians (50°11’26” N, 5°10’51” W) and site *Z*, a farm approximately 1 km away (50°11’56” N, 5°10’9” W) (Figure 1a). Jackdaws at the study sites are captured and colour-ringed for individual identification, and one of the colour rings contains a uniquely identifiable radio frequency identification *(RFID)* tag (*IB Technologies*, UK). Pair bonds and first-order kin relationships were determined through long-term monitoring of sightings, nest-box occupancy, and breeding attempts.

**Figure 1.**
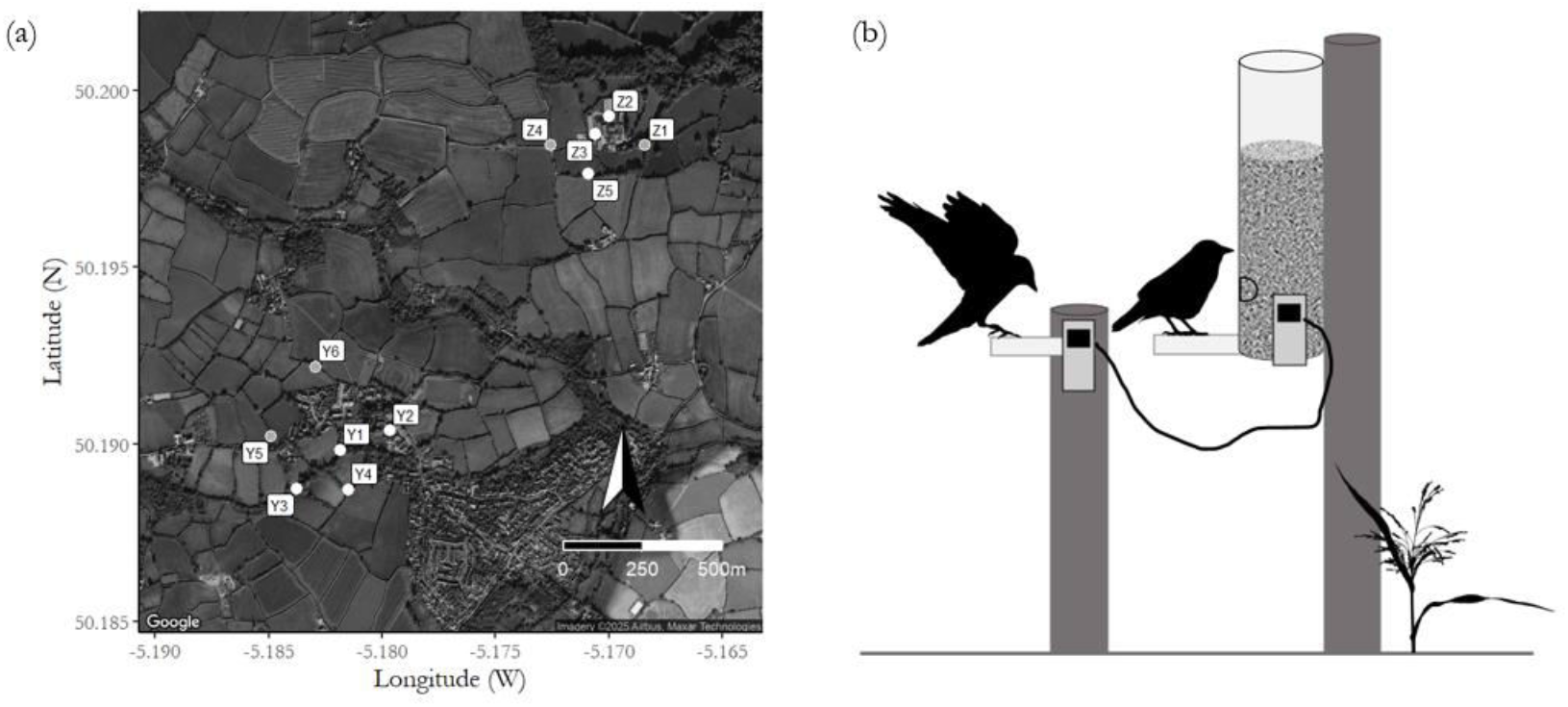
*(a)* Locations of feeding stations at the two study sites *Y* and *Z*. White circles indicate dyadic feeding stations with two *RFID* perches, and grey circles indicate stations with a single perch. At *Y2* we deployed an additional secondary perch towards the end of the data collection period. The feeding station at *Y5* had to be relocated to *Y6. (b)* Setup of dyadic feeding stations with primary (feeder) and secondary perches (note that *RFID* feeding station and logger design are simplified representations). At single-perch feeding stations, the secondary perch behind the primary feeding perch was not present. Jackdaw silhouettes were created by Josh Arbon, and the grass silhouette was taken from *PhyloPic* (by T. Michael Keesey).

### Data collection

During the 2023 breeding season (18 March to 04 August), we deployed *RFID* feeding stations (*Nature Counters Ltd*, UK) at 11 different locations across the two sites (six at site *Y*, five at *Z*; Figure 1a). Feeding stations contained an *RFID* antenna that detected the identity of visiting *RFID*-tagged jackdaws, as well as the date and time of their visit. Feeders were approximately 1 m above the ground, and provided a mixture of seeds, grain, and mealworms *ad libitum* to tagged jackdaws. Crucially, six of the locations were “dyadic” feeding stations, with an adjacent secondary *RFID* perch approximately 20 cm behind the main feeder (Figure 1b). Here, one jackdaw could feed while another “queued” on the perch immediately behind the feeding individual, allowing us to quantify levels of social tolerance. Feeding (primary) and queuing (secondary) perches were synchronised by a network cable allowing us to detect “dyadic events”, where two birds were detected at both perches simultaneously. We refer to the bird arriving at one of the two perches first as the “initiator”, and the second bird as the “joiner”. In 88% of dyadic events, the initiator sat on the feeder perch and the joiner on the secondary perch; for the other 12% of events the positions were reversed. Single-perch feeding stations (i.e. without a secondary perch) served to collect additional data on individuals’ propensity to use feeders, as well as on general social associations. During the study period (139 days; ∼ 20 weeks), *RFID* feeding stations recorded visits by 460 jackdaws (133 females, 169 males, 158 unsexed; 114 of the unsexed birds were juveniles that fledged during the study). Of these, 260 individuals (71 females, 100 males, 89 unsexed) were involved in at least one dyadic event. We recorded 2,703 dyadic events in total, of which 2,515 events (93%) occurred in the period after the first chicks had fledged. Dyadic events involved 1,001 unique dyads (943 during the period in which juveniles were present), 23 and 61 of which had been identified as pairs and first-order kin, respectively. Over the course of the data collection period, we detected 114 juveniles at *RFID* feeding stations, of which 65 participated in at least one dyadic event. 74 juveniles were, at some point, detected at primary perches of dyadic feeding stations (with feeding station *Y2* considered as a single-perch station in this analysis given that it was set up as a dyadic station later in the study period). Of those 74 juveniles, (24 “Dyadic” juveniles queued behind their parents at some point and 50 “Solo” juveniles did not queue behind their parents).

### Statistical analyses

Data were processed and analysed in *R* version 4.3.2. (47) via *R studio* version 2023.12.0 (48). We constructed (*Generalised) Linear Mixed Models ((G)LMMs)* using the package *glmmTMB* (49). We used the packages *DHARMa* (50) for model diagnostics and *car* (51) to test for collinearity using the variance inflation factor *(VIF)* and to conduct statistical inference by obtaining *Wald χ*^*2*^ statistics. We conducted pairwise comparisons of groups within a statistically significant categorical variable via the package *emmeans* (52). We plotted the data and model outputs using the packages *ggplot2* (53) and *sjPlot* (54). In most statistical models, the main independent variable of interest was the relationship type between two birds, classified into three categories: “Pair” (mated partners that had previously been identified through observation), “Kin” (first-order kin, i.e. parents and offspring, as well as siblings, previously identified by monitoring breeding attempts at nestboxes), or “Other” (non-kin and/or non-pairs, or no information about relationship type available). In all models (unless mentioned otherwise), we fitted “initiator ID”, “joiner ID”, and “dyad ID” as random effects to account for individual and dyad-level variation and dependencies, and for pseudo-replication. We also fitted “feeding station ID” as a random effect to account for potential variation across locations. We present key results in the main text and refer to the Supplementary Material for full model outputs. To investigate to what extent dyads were consistent (i.e., repeatable) in (i) their coordination (see *(2) coordination* below) and (ii) their social tolerance (see *(3) social tolerance* below) during foraging, we quantified the repeatability of the corresponding variables of interest, using a mixed effects modelling approach via the package *rptR* (55). We quantified repeatability at the level of unique dyads and included the type of their relationship to obtain an adjusted repeatability. We set the number of parametric bootstrapping and permutations to 1000.

#### (1) Social bias

To test the prediction that jackdaws preferentially forage with bonded partners, we constructed *GLMM1* (using a *Conway-Maxwell* distribution to account for underdispersion) with the “number of dyadic events per dyad” as a dependent variable and “relationship type” as an independent variable. To account for potential biases and assortment based on age class (56), we added the age class combination of both birds irrespective of the order of arrival (“adult-adult”, “adult-juvenile”, “juvenile-juvenile”). In this model, the IDs of the two individuals were included separately, but “dyad ID” was not included as a random effect because only one observation (number of dyadic events throughout the study) per dyad was used. To enable comparisons across relationship types, we only used data from the period in which juveniles started using feeders after fledging (i.e., 2515 out of 2703 dyadic events). To visualise how often birds engage in dyadic events we used the package *igraph* (57) to construct a weighted social network based on frequency of dyadic events detected at six dyadic feeding stations. Edges corresponded to instances of dyadic events between two individuals, with the direction indicating who queued behind whom (with the arrow pointing from the queuing to the feeding individual; see Figure 2c).

**Figure 2.**
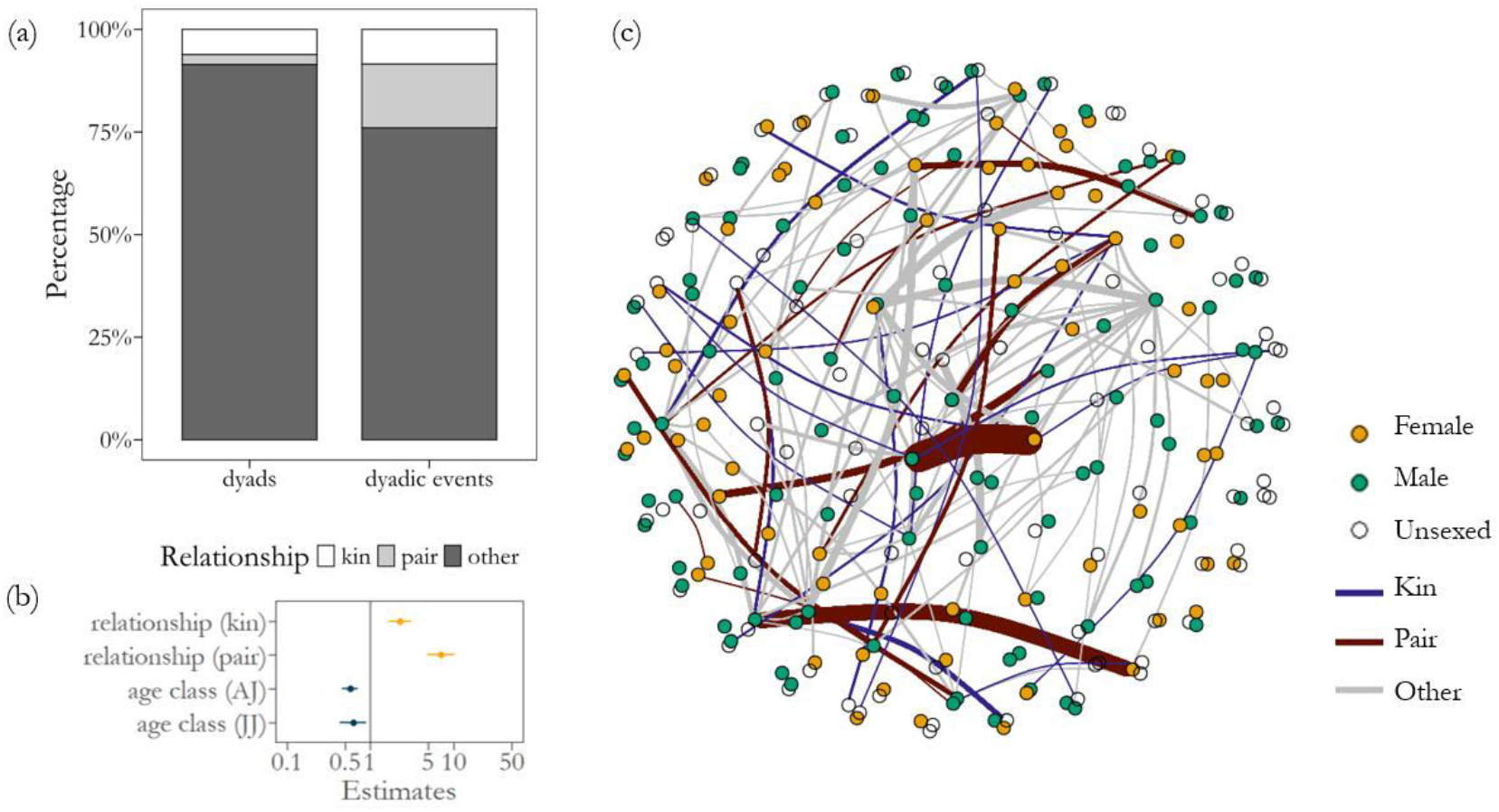
*(a)* Percentage of dyads (out of N = 943) and dyadic events (out of N = 2515 dyadic events) per relationship type. *(b)* Model estimates (dots) and standard errors (horizontal lines). *(c)* Weighted social network of dyadic events between individuals (i.e., proximity network). For visualisation purposes, the network is thresholded to include edges between dyads that queued together at least three times. The thickness of edges reflects the frequency with which individuals queued behind others.

#### (2) Coordination

To test our prediction *(a)* that bonded partners show greater temporal coordination, we examined the latency to arrive at feeding stations between both birds during dyadic events. We constructed *GLMM2* (*Gamma* family to account for the skewed distribution of continuous data) with the “latency” (the difference in the arrival time of the two birds per dyadic event (s)) as a dependent variable and the “relationship type” as an independent variable. The “age class combination” of both birds irrespective of the order of arrival (“adult-adult”, “adult-juvenile”, “juvenile-juvenile”) was fitted as an additional independent variable.

#### (3) Social tolerance, benefits, and costs

To test the prediction *(a)* that bonded partners perch together for longer (i.e. show greater social tolerance) we constructed *GLMM3* (*Gamma* family to account for the skewed distribution of continuous data) with the duration per dyadic event (s) as a dependent variable, and “relationship type” as the key independent variable. To account for variation associated with differences in competitive ability, which may partly be based on age, we also included “age class combination”. As the degree of familiarity between individuals could also influence the duration of dyadic events, we fitted the “number of dyadic events per dyad” indicative of familiarity. The number of dyadic events per dyad in this context was positively correlated with the dyadic association strength in an association network generated from both solo and dual feeders (see Supplementary Material for how this association network was constructed). To control for inter-individual variation in the propensity to use feeding stations in general we also added “initiator visit number” and “joiner visit number” (total visit number at all feeding stations, including single-perch feeding stations)

To test the prediction *(b)* that individuals would feed for longer with bonded partners than when with non-bonded birds we constructed *GLMM4* (*Gamma* family), which considered the total duration of feeding bouts (this differs from *GLMM3* as it includes the time that feeding individuals spent at the feeder perch *before or after* the dyadic event occurred, and it included instances in which the initiator sat at the feeder perch). We fitted the “time that the initiator spent at the feeder perch” (as a proxy for food intake; see Supplementary Material for validation of food intake based on visit duration) as a dependent variable and with the same independent variables as mentioned above. In these two models, we additionally controlled for the latency of arrival time between both birds. Numeric variables were scaled (z-transformed) for enhanced model performance and comparability in *GLMM3* and *GLMM4*.

Finally, following from *GLMM4*, we constructed two GLMMs (*Gamma* family) to examine the averaged visit duration at feeding perches by paired adults (N = 32; *GLMM5*), and juveniles (N = 10; *GLMM6*) that visited feeders in all of three different social contexts: (i) with a bonded adult queuing behind (pair-bonded mate in the case of adults, and parents in the case of juveniles), (ii) another non-bonded adult queuing behind, and (iii) no individual queuing behind. We report results from models including averaged individual-level visit duration as we were interested in individual-level comparisons, but the results were qualitatively equivalent to visit-level analyses.

#### (4) Social learning

To test whether juveniles learned socially about locations of feeding stations by queueing behind their parents (prediction *(a)*), we compared juveniles that joined their parents at feeders (“Dyadic” juveniles) with juveniles that did not (“Solo” juveniles; note that these juveniles may still have associated with their parents outside of the experimental context). Specifically, we examined whether dyadic juveniles made their first independent visit to feeding (primary) perches of dyadic feeding stations earlier than Solo juveniles (dependent variable 1, “first visit, numeric day in the study period”; excluding position *Y2* from the analysis because it was deployed later in the study period). We also tested whether dyadic juveniles visited feeding perches of dyadic feeding stations more often than Solo juveniles, considering different periods (dependent variable 2, “number of visits per juvenile per period”). If they learned from parents, then dyadic juveniles would be expected to visit feeders sooner (dependent variable 1) and more often (dependent variable 2) than Solo juveniles. We selected periods for comparison based on the average day at which juveniles were first detected engaging in dyadic events with their parents (“mean first day”, i.e., study day 108). From that day, we selected two periods of two weeks each before and after the “mean first day”, and also periods of more than two weeks before and after that day (16 and 19 days long, respectively). We fitted one (*G)LMM* for each dependent variable (“study day of first visit”, *LMM7*; “number of visits”, *GLMM8, Poisson* family for count data), with an interaction between juvenile group (“dyadic”, “solo”), and phase (“more than 2 weeks before”, “2 weeks before”, “mean first day”, “2 weeks after”, “more than 2 weeks after”) as independent variables. We fitted “juvenile ID” as a random effect to account for repeated measures per juvenile across the different periods. To examine formally whether dyadic juveniles were overrepresented in terms of their visit number, we also conducted a Chi-square test. Finally, to test prediction *(b)*, we repeated the analysis equivalent to *GLMM8* to test whether naïve juveniles who had not followed their parents (Solo juveniles) may have learned about feeding stations by queuing behind knowledgeable juveniles (Dyadic juveniles) who had previously associated with their parents (“number of visits”, *GLMM9*).

## Results

### (1) Social bias

As predicted, jackdaws were more likely to engage in dyadic events with bonded partners than with others. The number of dyadic events differed according to relationship type (*GLMM1, relationship type, χ*^*2*^ = 402.102, df = 2, *P* < 0.001) (Figure 2; Supplementary Material, Table S1). Pair-bonded mates engaged in more dyadic events (mean ± sd = 18.09 ± 28.93 absolute visits per dyad) than kin (3.67 ± 3.50 visits per dyad) (*kin-pair, estimate* = - 1.419, *SE* = 0.165, *z* = - 8.581, *P* < 0.001) and “others” (2.33 ± 3.36 visits per dyad) (*other-pair, estimate* = - 2.224, *SE* = 0.115, *z* = - 19.271, *P* < 0.001). Kin also participated in more dyadic events than other dyads (*other-kin, estimate* = - 0.805, *SE* = 0.124, *z* = - 6.476, *P* < 0.001). Approximately 82% of dyadic events between kin were between parents and their offspring, and the remaining events occurred between siblings. The number of dyadic events also differed according to age class combination (*GLMM1, age class combination, χ*^*2*^ = 8.046, *df* = 2, *P* = 0.018) (Figure 2b; Supplementary Material, Table S1).

### (2) Coordination

#### (a) Latency of arrival

As predicted, bonded partners (pairs, kin) were more coordinated during foraging (*GLMM2, relationship type, χ*^*2*^= 39.444, *df* = 2, *P* < 0.001; Figure 3a, b; Supplementary Material, Table S2). Arrival latency was significantly, and nearly 40% shorter for “kin” (18.33 ± 19.45 s) (*other-kin, estimate* = 0.366, *SE* = 0.088, *z* = 4.172, *P* < 0.001) and “pairs” (19.99 ± 26.19 s) (*other-pair, estimate* = 0.411, *SE* = 0.084, *z* = 4.876, *P* < 0.001) compared to “other” dyads (30.85 ± 31.21 s). By contrast, arrival latencies between “kin” and “pair” dyads did not differ significantly (*kin-pair, estimate* = 0.045, *SE* = 0.119, *z* = 0.378, *P* = 0.924). Age class combination was not associated with the latency to arrive (*GLMM2, age class combination, χ*^*2*^= 3.094, *df* =2, *P* = 0.213) (Figure 3b; Supplementary Material, Table S2).

**Figure 3.**
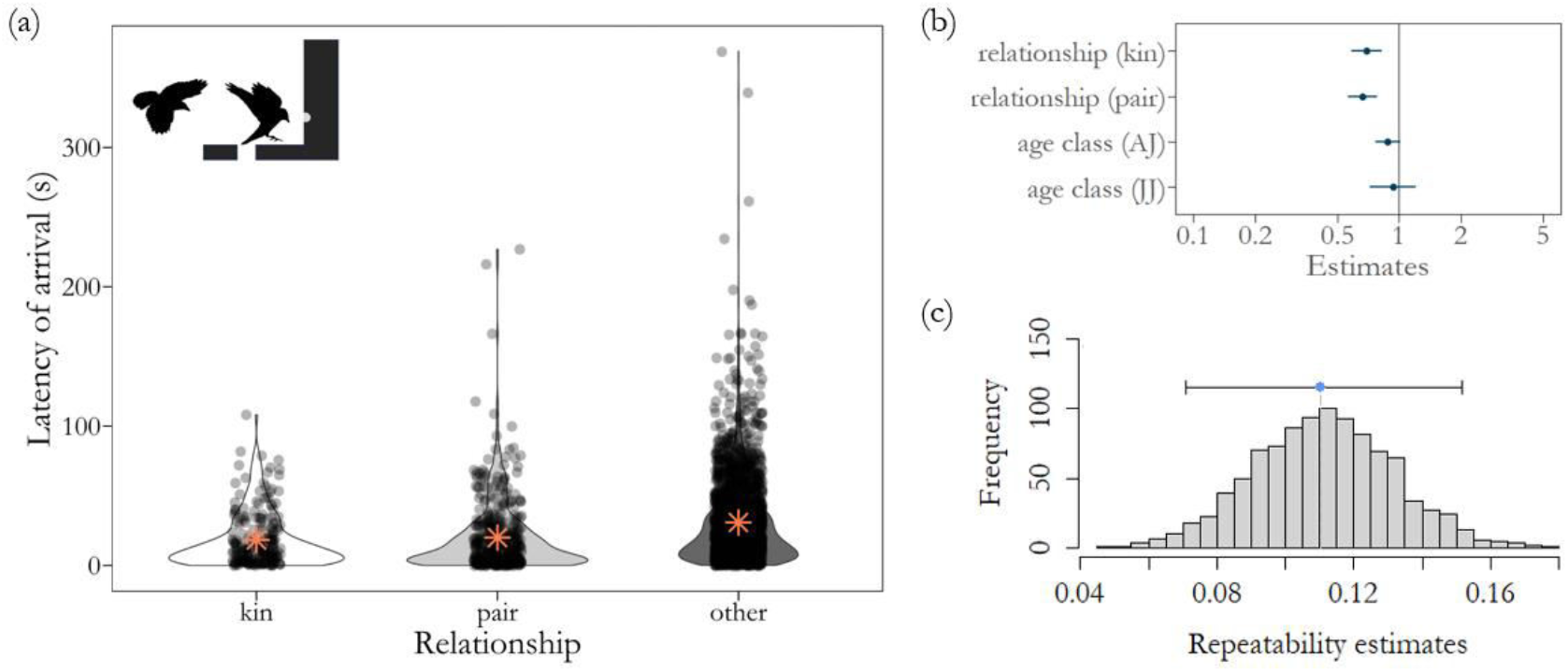
(a) Latency of arrival between two individuals engaging in dyadic events (each datapoint reflects a single dyadic event; red asterisks show means per relationship type). (b) Model estimates and errors for two dependent variables, “relationship” and “age class combination”. (c) Repeatability of latencies (blue point = repeatability estimate; horizontal bar = 95% confidence interval). The jitter-position parameter and the alpha parameter adjusting transparency of data points based on density were set to 0.1 and 0.3, respectively. Jackdaw silhouettes by Josh Arbon.

#### (b) Repeatability

The latency in arrival time was repeatable for dyads, accounting for relationship types (*R*_*adjust*_ = 0.111, *SE* = 0.021, *CI* = [0.071, 0.156], *PLRT* < 0.001, *Pperm* = 0.001) (Figure 3c).

### (3) Social tolerance

#### (a) Social tolerance and opportunity cost

Dyadic events lasted longer between bonded partners than between others, with the queuing individual incurring a corresponding opportunity cost. On average, events lasted around three times longer between pair partners 12.28 ± 13.85 s (mean ± sd) and between first-order kin (10.41 ± 11.50 s) compared to other dyads (3.81 ± 5.93 s) (*GLMM3, relationship type, χ*^*2*^ = 92.593, *df* =2, *P* < 0.001; Figure 4a, c, d; Supplementary Material, Table S3). Posthoc comparisons indicated that dyadic events lasted longer for pairs than for “other” dyads (*other-pair, estimate* = - 0.965, *SE* = 0.107, *z* = - 9.030, *P* < 0.001) and for kin (*kin-pair, estimate* = - 0.638, *SE* = 0.132, *z* = - 4.855, *P* < 0.001), and longer for kin than for “other” dyads (*other-kin, estimate* = - 0.327, *SE* = 0.084, *z* = - 3.904, *P* < 0.001). The duration of dyadic events was longer for dyads that engaged in these visits more frequently (and were potentially more familiar), controlling for individual propensity of initiators and joiners to use feeding stations (*GLMM3, visit number dyad, β* = 0.163, *SE* = 0.064, *χ*^*2*^ = 6.429, *df* = 1, *P* = 0.011; Figure 4c; Supplementary Material, Table S3). This result remained when excluding bonded individuals, i.e. kin and pairs (*GLMM3, visit number dyad, β* = 0.100, *SE* = 0.040, *χ*^*2*^ = 6.356, *df* = 1, *P* = 0.012), and a non-significant trend remained when excluding putative pairs (mixed-sex dyads whose pair-status was uncertain) (*GLMM3, visit number dyad, β* = 0.074, *SE* = 0.039, *χ*^*2*^ = 3.545, *df* = 1, *P* = 0.059). The duration of dyadic events was also linked to the age class combination of both individuals (*GLMM3, age class combination, χ*^*2*^ = 72.464, *df* =2, P < 0.001; Figure 4c; Supplementary Material, Table S3). For full model output see Supplementary Material (Table S3).

**Figure 4.**
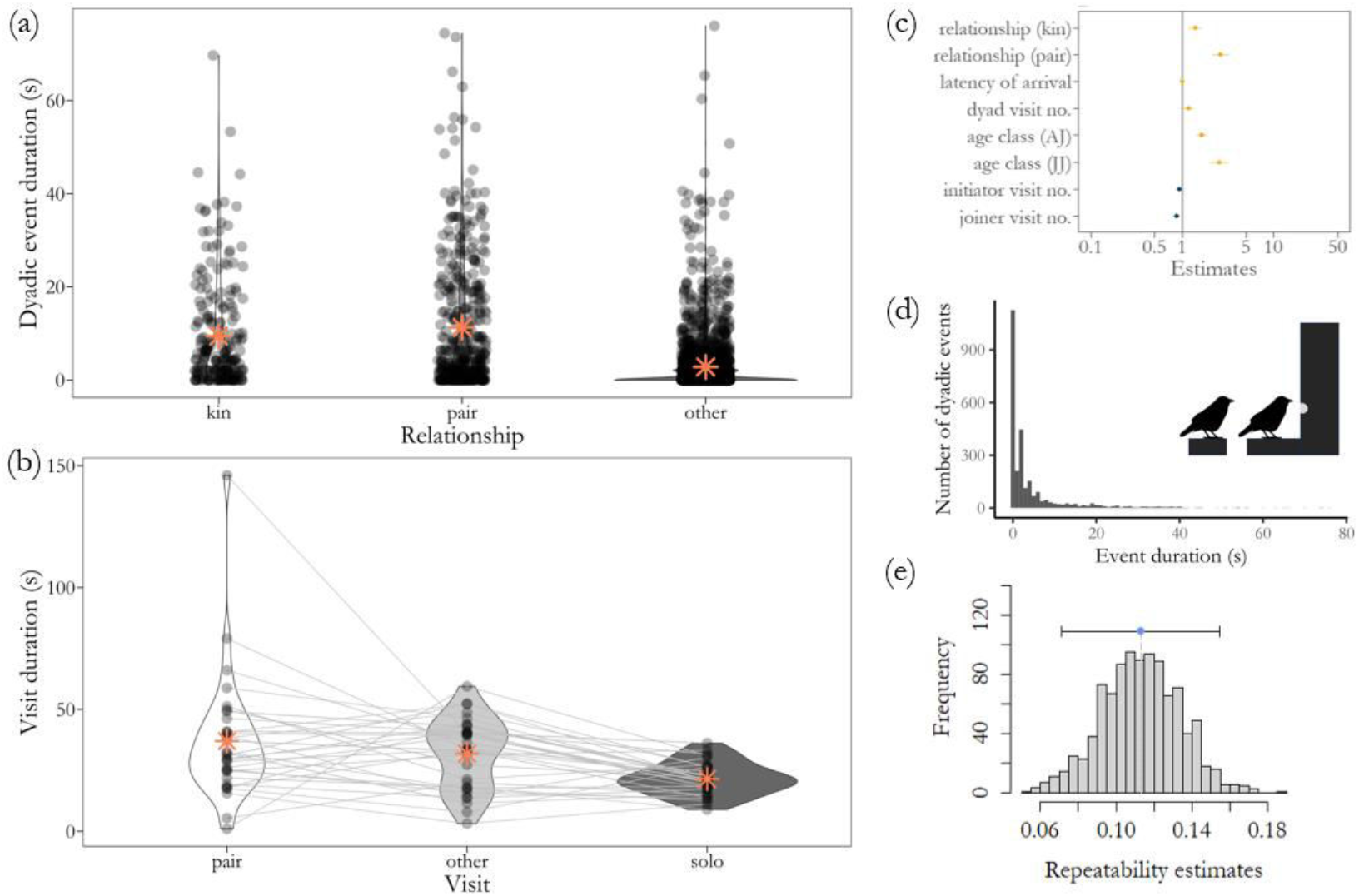
*(a)* Duration of dyadic events as a function of the relationship type between different birds. The jitter-position parameter and the alpha parameter adjusting transparency of data points based on density were set to 0.1 and 0.3. *(b)* Average visit duration of N = 32 paired individuals at the primary feeder perch when accompanied by their pair-bonded mate (“pair”), when accompanied by another adult (“other”) or when alone (“solo”). *(c)* Model estimates from *GLMM3* (data shown in *(a)*).

#### (b) Forager benefit

The presence of bonded individuals provided foraging benefits for the feeding individual. Individuals perched at the feeding (primary) perch could feed for longer if their mate was on the queuing perch behind them (see Supplementary Material, “Validation of food intake” and Figure S1 for validation that visit duration at *RFID* feeding stations corresponded to feeding bouts, i.e., likely food intake). The time spent feeding differed according to relationship type (*GLMM4, relationship type, χ*^*2*^ = 42.002, *df* =2, *P* < 0.001; see Table S4 for full model output) with the initiator accompanied by their mated partner staying for longer than those accompanied by kin (*kin-pair, estimate* = - 0.340, *SE* = 0.079, *z* = - 4.333, *P* < 0.001) or “other” individuals (*other-pair, estimate* = - 0.382, *SE* = 0.059, *z* = - 6.475, *P* < 0.001), but there was no difference between kin and “other” dyads (*other-kin estimate* = - 0.042, *SE* = 0.056, *z* = - 0.741, *P* = 0.739).

Analysis of the subset of paired birds and juveniles that visited feeders in three different contexts (together with a bonded adult partner, another adult, or alone) showed evidence that individuals obtained benefits by feeding for longer if their partner or parent, respectively, was queuing behind them than when they were accompanied by another adult or when they were alone (Figure 4b; Supplementary Material; Table S5, Table S6). Visit duration of paired individuals at primary perches was associated with the social context (*GLMM5, visit type (paired mate vs another adult vs solo), χ*^*2*^= 53.322, *df* =2, *P* < 0.001; Figure 5b; Table S5). Paired birds stayed at feeders for longer when accompanied by their mate compared to when another adult was present (*pair-other, estimate* = 0.404, *SE* = 0.094, *z* = 4.309, *P* < 0.001) and compared to when they were alone (*pair-solo, estimate* = 0.663, *SE* = 0.091, *z* = 7.301, *P* < 0.001). Furthermore, birds stayed at feeders for longer when another adult was queuing behind compared to when they were alone (*other-solo, estimate* = 0.259, *SE* = 0.085, *z* = 3.045, *P* = 0.007). Similarly, visit duration of juveniles at primary perches was associated with the social context (*GLMM6, visit type (parents vs another adult vs alone), χ*^*2*^= 36.572, *df* =2, *P* < 0.001; Supplementary Material, Table S6, Figure S2). Juveniles were perched at feeders for longer when their parent queued behind them than when another adult was present (*parent-other, estimate* = 0.281, *SE* = 0.098, *z* = 2.856, *P* = 0.013) and compared to when they were alone (*parent-solo, estimate* = 0.593, *SE* = 0.098, *z* = 6.044, *P* < 0.001). Juveniles also stayed at feeders for longer when accompanied by another adult than when they visited alone (*other-solo, estimate* = 0.312, *SE* = 0.098, *z* = 3.183, *P* = 0.004).

**Figure 5.**
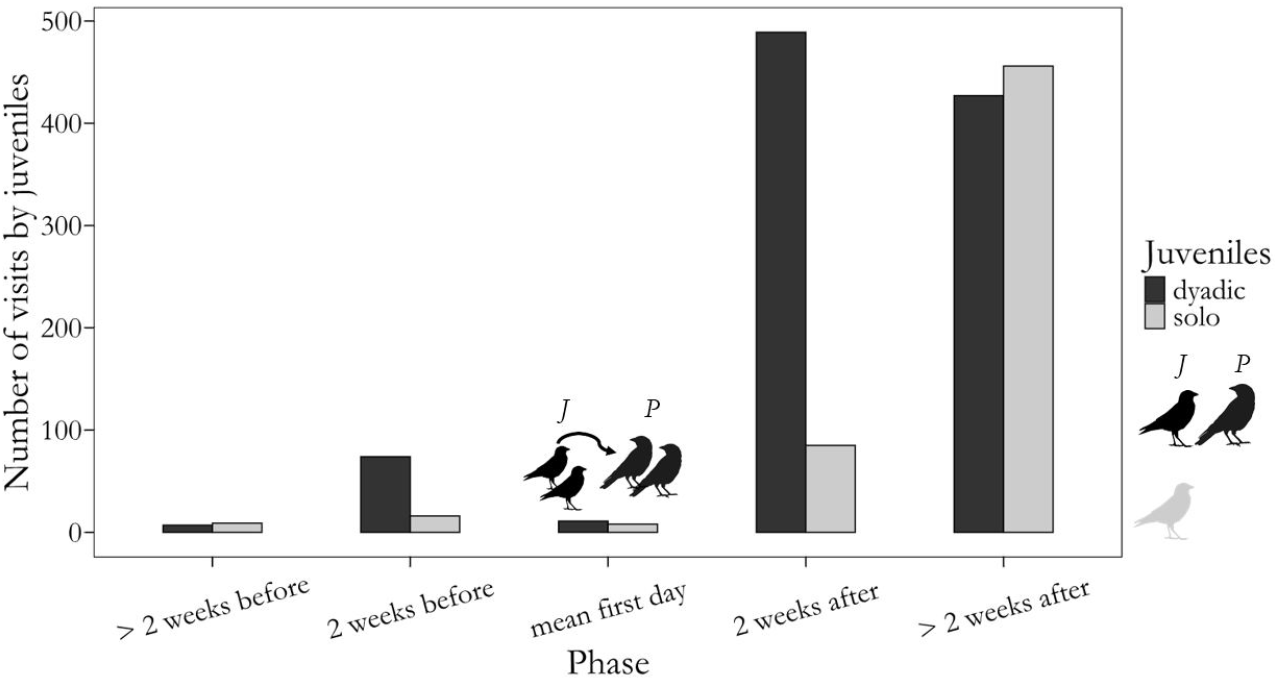
Total number of independent visits made by juveniles during different phases. The number of visits by juveniles of two “groups” (Dyadic, N = 24 juveniles *(J)* engaging in dyadic events with their parents *(P)*, and Solo, N = 50 juveniles not engaging in dyadic events with their parents) is depicted in this bar chart for different phases. The phase “mean first day” denotes the average day of the study period on which juveniles of the Dyadic group first engaged in dyadic events with their parents (day 108 of the study). The other phases were divided into 2-week intervals to compare changes in visit rates between the two groups. Jackdaw silhouettes were obtained from *PhyloPic* (uploaded by Birgit Lang and Ferran Sayol).

#### (c) Repeatability

The duration of dyadic events was weakly and significantly repeatable for dyads, accounting for relationship types (*R*_*adjust*_ = 0.113, *SE* = 0.021, *CI* = [0.071, 0.154], *PLRT* < 0.001, *Pperm* = 0.001; Figure 4e). *(d)* Distribution of the durations of dyadic events (s) *(e)* Repeatability plot for the duration of dyadic events per dyad (blue point = repeatability estimate; horizontal bar = 95% confidence interval). Jackdaw silhouettes were created by Josh Arbon.

### (4) Social learning

### (a) Vertical social learning

Engaging in dyadic events with parents appeared to promote social learning of feeder usage by juveniles. Juveniles that engaged in dyadic events with their parents went on to visit feeders independently (i.e., without their parents) earlier and more often than juveniles who did not queue behind their parents. Although only a minority of juveniles (∼32%) engaged in dyadic events with their parents, they accounted for 1008 out of 1582 (∼64%) of independent visits made by juveniles at primary perches of dyadic feeding stations, (Chi-square test, *χ*^*2*^ = 28.147, df = 1, *P* < 0.001). Dyadic juveniles made their first independent visit to *RFID* feeding stations significantly earlier than Solo juveniles (*LMM7, juvenile group, β* = 10.660, *SE* = 2.500, *χ*^*2*^ = 18.186, *df* = 1, *P* < 0.001; Supplementary Material, Table S7). On average, Dyadic juveniles made their first independent visit on the same day at which the first dyadic event with their parents occurred. Dyadic juveniles visited dyadic feeding stations around four times more often than Solo juveniles that had never queued behind their parents (mean ± sd = 42.00 ± 51.46 visits per Dyadic juvenile and 11.48 ± 20.85 per Solo juvenile from fledging until the end of the study period). The difference between Dyadic and Solo juveniles emerged particularly in the period after juveniles had first visited with their parents. This was evidenced by a significant interaction between juvenile group and phase (*GLMM8 phase * juvenile group, χ*^*2*^ = 198.130, *df* = 4, *P* < 0.001; Figure 5; Supplementary Material, Table S8).

### (b) Horizontal social learning

Solo juveniles that queued behind Dyadic juveniles who had previously queued behind their parents visited feeding stations more often. The difference in visit number between two groups of Solo juveniles (those who queued behind Dyadic juveniles and those who did not) arose particularly in the last phase, when Dyadic juveniles had already been at feeders with their parents: there was a significant interaction between the phase and the group (*GLMM9, phase * juvenile group, χ*^*2*^ = 13.062, *df* = 1, *P* < 0.001; Supplementary Material, Table S9).

## Discussion

We tested the overarching hypothesis that jackdaw social relationships shape decisions and interactions at monopolisable feeding stations, with consequences for the benefits and costs of acquiring food and information. As predicted, we found that visits to feeding stations by bonded partners were more frequent, better coordinated, and longer. This meant that compared to other pairs, bonded partners had greater foraging intake (for the first arriving bird), and social learning opportunities (for juveniles), but short-term opportunity costs (for the bird queuing).

Bonded partners visited feeding stations together more frequently (nine times more often in the case of pair-bonded mates) and arrived with shorter latencies (around 40%) compared to other dyads. The time that elapsed between the first arriving bird and the partner was also repeatable within dyads, which suggests coordination of foraging. Being regularly accompanied by, and coordinating foraging activities with, bonded partners can make foraging outcomes at monopolisable resources more predictable, for example in terms of accessing and defending resources. This is particularly relevant in a social system that is dominated by dynamic, potentially unpredictable, and competitive social interactions, as observed in jackdaws (56). By coordinating foraging activities, individuals may be reliably accompanied by a social partner that could provide social support (21). Such coordination may also reduce the cost of searching for social partners, or other costs associated with social foraging, such as competition (58).

We find that jackdaws in bonded pairs are willing to incur opportunity costs, foregoing foraging to queue behind social partners. Similar findings have been reported in studies on other monogamous social corvids. In rooks *(Corvus frugilegus)*, high levels of social tolerance within mated pairs facilitated access to food for scrounging individuals (59) and in Northwestern crows *(Corvus caurinus)*, individuals were less aggressive towards close relatives (60). In non-breeder groups of common ravens, bonded individuals competed for food more successfully even in cases in which their partner was not immediately present (61). In our study, bonded partners obtained immediate foraging-related benefits by increasing feeding duration (and thus food intake) through social tolerance. Such short-term foraging benefits may lead to longer-term fitness benefits, such as better body condition and health, and potentially greater survival and/or reproductive success (e.g., (62)). This could be particularly relevant for juveniles who lack competitive ability and need to ensure consistent food intake over the course of their development, and we expect that greater social tolerance among kin and mated partners provides indirect fitness benefits due to kin selection (63) and high fitness interdependence between genetically monogamous mated partners (37,64), respectively. Future work could investigate the temporal dynamics of queuing in our system; for example, if and how tolerance changes with variable foraging payoffs (65–67) and whether individuals might alternate roles across interactions, taking turns to feed and queue in a form of reciprocal cooperation. Such turn-taking could potentially help stabilise cooperative partnerships (68) reinforcing the mutual benefits of maintaining close social bonds.

Our findings also highlight a potential role for familiarity in social foraging interactions. We found that dyads that visited more often together also tolerated each other for longer, even when excluding bonded individuals (pairs and kin) from the analysis (and controlling for individual variation in feeder use). This indicates that social tolerance may be afforded to frequent or recent associates that are not pair-bonded mates or kin, i.e. “friends” (2,24). Such encounters by familiar individuals can be characterised by relatively higher levels of social tolerance as compared to transient acquaintances and strangers. Jackdaws encountering each other frequently at feeding stations might have learned to tolerate each other more over time (42) or could have already been associating together previously, outside of this experimental context. Similarly, the strength of previous associations predicted social tolerance at food patches in common marmosets *(Callithrix jacchus)* (28). Further research is needed to better understand the role of familiarity in non-pair and non-kin relationships during social foraging in jackdaws and other species.

We also found that, as predicted, social tolerance between parents and offspring promoted social learning about feeding stations by juveniles in a phase during which they were likely naïve about the distribution of food resources. Juvenile jackdaws that associated closely with queued behind their parents by queuing behind them (Dyadic juveniles) visited feeding stations independently earlier and more often compared to juveniles that did not engage in dyadic events with their parents (Solo juveniles, who may nonetheless have associated with their parents outside of the particular experimental context). This difference in visit rates between Dyadic and Solo juveniles was particularly evident in the two weeks after Dyadic juveniles started queuing behind their parents, suggesting vertical transmission of information about feeding stations. Dyadic juveniles also seemed to visit feeding stations more often than Solo juveniles even in the two weeks before the average day at which they first queued behind their parents. This is probably because the first juveniles had already started queueing behind their parents during those two weeks and had the opportunity to make use of the new knowledge. We cannot rule out the possibility that Dyadic juveniles were, simply by chance, more likely to visit feeding stations irrespective of dyadic events with their parents serving as social learning opportunities. However, this is unlikely because on average, Dyadic juveniles made their first independent visit on the same day that they queued behind their parents. Social learning early in life may have long-lasting consequences, as seen in blue tits *(Cyanistes caeruleus)* and great tits, where foraging behaviour that was socially learned from foster parents persisted throughout life (69). In long-lived species with long periods of parental care, such as corvids, vertical cultural transmission of social information may be particularly valuable (70). Moreover, individuals that learn from parents may then serve as sources of information for their peers, enabling horizontal transmission. This was seen in our study, where Dyadic juveniles are likely to have served models for other juveniles because juveniles tend to associate with each other after fledging (56). Among Solo juveniles (who never queued behind their parents), those who then queued behind other juveniles visited feeding stations more often than those who did not. This effect particularly emerged in the period after juveniles had become knowledgeable by following their parents. Thus, juvenile birds may partly use social information from both their parents and their peers. For example, in juvenile hihis *(Notiomystis cincta)*, early foraging decisions were influenced by their parents and later on by their peers (71). Obtaining information by associating with more knowledgeable individuals could be one of the underlying causal mechanisms of the apparent link between sociality and long-term fitness.

Overall, we showed that close, long-term social relationships shape the likelihood and outcomes of fine-scale dyadic foraging interactions in a dynamic social system of wild jackdaws. In this social system, characterised by high levels of fission-fusion dynamics (44), individuals may be faced with high levels of social uncertainty and competition due to the variable and transient nature of social interactions with a high number of different other individuals. Coordinating with and tolerating bonded social partners with shared interests can enable individuals to manage uncertainty and increase short-term access to food and information. Thus, through their daily social decisions, individuals may construct their own immediate social niches (72) to balance the benefits and costs of sociality and maximise short-term payoffs, with important effects on long-term fitness.

## Ethics statement

Ethical approval for this study was granted by the *University of Exeter Biosciences Research Ethics Committee* (512510). Researchers adhered to the *ASAB* guidelines for the ethical treatment of nonhuman animals in behavioural research and teaching (73). Researchers captured and ringed birds and collected blood samples under licences from the *British Trust for Ornithology (BTO)* and *UK Home Office* (project licence P882CF514).

## Data and code

Data and code will be available online […].

## Conflicts of interest

The authors have no conflicts of interest to declare

## Author contributions

LGH – conceptualisation, data curation, formal analysis, investigation, methodology, project administration, software, validation, visualisation, writing – original draft, writing – review and editing

GEM – data curation, project administration, resources, writing – review and editing

IF – supervision, writing – review and editing

AJK – supervision, writing – review and editing

AT – conceptualisation, funding acquisition, methodology, project administration, resources, supervision, writing – review and editing

## Funding

L.G.H. was supported by the Biotechnology and Biological Sciences Research Council-funded South West Biosciences Doctoral Training Partnership (DTP3: BB/T008741/1). A.T. was supported by a Leverhulme Trust grant (RGP-2020-170).

## Acknowledgements

We are grateful to people in Stithians, particularly Odette Eddy, as well as the Gluyas family and staff at Pencoose Farm for granting access to their land to conduct this research. Many thanks to Josh Arbon for providing initial guidance for the use of *RFID* equipment, for discussion, and for sharing jackdaw silhouettes. We also thank Andrew Young for additional discussion. Thanks to Zazie Benoit-Delaby for coding videos for validation of feeding bouts, and to team members of the *Cornish Jackdaw Project* for occasionally assisting in the field.

## Notes

### Competing Interest Statement

The authors have declared no competing interest.

